# Spinal cord Ca^2+^ imaging reveals glial-driven central sensitization in post-traumatic osteoarthritis

**DOI:** 10.64898/2026.06.01.729291

**Authors:** Samuel W. Fung, Erika K. Harding, Jenny K. Cheung, Hantao Zhang, Julieanne L. T. Dalsgaard, Stephanie M. Norlock, Jeff Biernaskie, John R. Matyas, Michael E. Hildebrand, Jo Anne Stratton, Robert P. Bonin

**Author notes:** These authors contributed equally (Samuel W. Fung, Erika K. Harding).

## Abstract

Central sensitization may be defined behaviourally, cellularly, or molecularly; yet these can all vary depending on the model and duration. Current electrophysiological approaches are time and labour intensive. Here, we developed a Ca^2+^ imaging and analysis pipeline (CuMIN) that implements semi-automated detection and analysis of cellular Ca^2+^ activity in rodent spinal cord slices, from which distinct signatures were defined for various acute and chronic pain models. Spinal cord slices from male mice were isolated after inducing pathological pain in a variety of well-established surgical or pharmacological approaches, incubated in a cell-permeant Ca^2+^ indicator, and imaged with epifluorescence microscopy. Intensity and temporal features of spontaneous and glutamate-evoked Ca^2+^ events were processed by linear discriminant analysis to map unique clusters of activity for each pain model. The resulting activity map of spinal dorsal horn activity is substantially different in the surgical model of chronic pain induced by post-traumatic osteoarthritis (PTOA), which lacks clear mechanistic evidence of central sensitization. Specifically, the PTOA Ca^2+^ activity signature overlapped with chemotherapy-induced neuropathy and neuropathic pain models, both of which are associated with gliosis-induced central sensitization. We confirmed gliosis in the PTOA model by immunostaining IBA1 and GFAP and observed analgesic effects of intrathecal carbenoxolone, Gap27, and minocycline that targeted glial activity. These findings validate CuMIN as a sensitive and specific approach for defining the basic cellular signatures of spinal central sensitization, with the utility of identifying potential therapeutic targets and serving as a translational platform for novel drug discovery across various acute and chronic pain models.

## 1. Introduction

Chronic pain is a heterogenous condition with differing etiologies.[45,73] Central sensitization, broadly defined by an increase in activity of central nociceptive neurons, is a critical driver of many chronic pain conditions,[16,91] yet it is unclear if central sensitization in different settings share similar or different pathophysiologies. Hence, it is of particular interest to compare and contrast the cellular pathophysiologies in various experimental chronic pain conditions to characterize cellular activity signatures of central sensitization, to define new targets and influence potential treatment options.

The mechanistic regulation of spinal central sensitization involves dynamic interactions among neurons, astrocytes, and microglia in the superficial dorsal horn (SDH) of the spinal cord.[16] At the neuronal level, SDH neurons exhibit enhanced excitability and activity through multiple mechanisms, including increased synaptic input, long-term potentiation at excitatory synapses, ion channel dysregulation, and disinhibition through reduced GABAergic and glycinergic transmission or KCC2 downregulation, such as during neuropathic pain.[31,47,63,82,91] Microglia can be activated early after peripheral injury and proliferate within the SDH, releasing pronociceptive mediators.[14,85] Astrocyte activation typically follows microglial activation and results in subsequent signaling that amplifies nociceptive transmission.[11,25,38] Importantly, the specific contributions of these cell types vary across pain conditions. In chemotherapy-induced peripheral neuropathy by paclitaxel, SDH astrocytic activation has been observed, with evidence of increased GFAP expression, and cytokine release.[78,98] In contrast, sciatic neuropathic pain models have a well-documented role of microglia in initiating and maintaining central sensitization.[14,39,63,84,85] In acute osteoarthritis models such as intra-articular monosodium iodoacetate (MIA) or collagenase injections, both SDH microglia and astrocytes are activated to varying degrees, with microglial P2X7 receptor signaling and pannexin-1 channels implicated in mechanical hypersensitivity.[1,61,64] Importantly, bidirectional signaling between neurons and glia creates feed-forward loops that can sustain pain states long after the initial peripheral insult has resolved.

Given the diverse and interconnected cellular and molecular mechanisms that contribute to pathological pain, there is a need for experimental approaches to assess changes in spinal network activity. Historically, central sensitization has been characterized using electrophysiological techniques, which are limited in terms of throughput. An ideal approach would study central changes in network activity with single-cell resolution. Therefore, we chose single-cell Ca^2+^ imaging of the SDH to evaluate local, *in-situ* activities of spinal nociceptive circuitry.

We developed an *ex-vivo* Ca^2+^ imaging analysis pipeline (CuMIN: **Cu**rated ROIs for **Min**ian) to assess central changes in pathological pain conditions. Using CuMIN, we analyzed SDH Ca^2+^ imaging data from different pathological pain models and discovered unique activity signatures of Ca^2+^ activity. We next investigated Ca^2+^ imaging data from spinal cords following post-traumatic osteoarthritis (PTOA), in which there is a lack of information about potential central mechanisms of central sensitization. We found that the signature of PTOA overlapped with those of SNI and paclitaxel, models that involve central sensitization associated with glial activation. Supporting this, we found that intrathecal injections of carbenoxolone, Gap27, and minocycline were analgesic for PTOA, indicating a role of spinal astrocytes and microglia in regulating mechanical hypersensitivity in PTOA and proposing a therapeutic avenue for this condition.

## 2. Methods

### 2.1 Animals

Animal handling and experimental procedures were performed at 2 locations: the University of Toronto and the University of Calgary. These procedures were conducted with the approval of the Local Animal Care Committees at their respective institutions at the University of Toronto and the University of Calgary, as well as the Canadian Council of Animal Care.

Experiments at the University of Toronto were performed with adult (>8 week old) male C57BL/6N mice purchased from Charles River (Sherbrooke, Quebec, Canada). Experiments at the University of Calgary were performed with adult male C57BL/6J mice, purchased from The Jackson Laboratory. Littermate controls were used whenever possible. Mice were bred and maintained on a fixed 14-hour light:10-hour dark (14:10 LD) cycle in groups of one to four mice per cage. Food and water were provided ad libitum.

### 2.2 Pain Models

#### Capsaicin Paw Injection

Mice were first lightly anesthetized with inhaled isoflurane (3%). Next, 1% w/v capsaicin (Capsaicin, Sigma-Aldrich, M2028) in a solution of 10:10:80 of ethanol:Tween-20:saline was injected into the plantar surface of one hind limb (5 μL) using a Hamilton syringe (Ethanol, House Brand, 39752-P016-EAAN; Tween-20; BioShop, TWN508.500). Mice were allowed to recover under observation and then utilized for experiments after 2 hours. Mechanical allodynia was observed as early as 1 hour after injection (data not shown). Contralateral non-injected tissue was used as controls.

#### CFA Paw Injection

Mice were first lightly anesthetized with inhaled isoflurane (3%). Next, Complete Freund’s Adjuvant (CFA, Sigma-Aldrich, F5881) was injected into the plantar surface of one forelimb (15 μL) using a Hamilton syringe. Mice were allowed to recover under observation and then utilized for experiments after 48 hours. Mechanical allodynia was observed as early as 3 hours after injection (data not shown). Contralateral non-injected tissue was used as controls.

#### SNI

Mice were first anesthetized (5% induction, 1.5% maintenance) with isoflurane, then a small incision was made on the skin of the left thigh. The muscle bundles were separated by blunt dissection to expose the three branches of the sciatic nerve (common peroneal, tibial, and sural nerves). The common peroneal and tibial nerves were tightly ligated with a 6/0 silk suture (Ethicon) and transected. A 1-mm piece of the nerve was removed to avoid regrowth. Precaution was taken to avoid damaging the sural nerve. After surgery, the overlaying muscles and skin were separately closed with 6/0 silk and 4/0 Vicryl sutures, respectively. After surgery, animals were allowed to recover for 14 days before experiments. Mechanical allodynia was observed as early as 1 week after surgery (data not shown). In sham-operated animals, the same procedure was performed but without ligation and transection of the nerves. Any mice that failed to demonstrate mechanical hypersensitivity after 14 days were excluded from future experiments.

#### Paclitaxel

Mice were first lightly anesthetized with inhaled isoflurane (3%). Paclitaxel (Selleck Chemicals, S1150) (6 mg/ml) in 50% El Kollipher (Sigma-Aldrich, C5135) and 50% ethanol was further diluted in sterile saline and administered via intrathecal injection at a dose of 50 nM in 5 μL saline. After injection, animals were allowed to recover for 2 hours before experiments. Mechanical allodynia was observed as early as 30 minutes after injection. Control animals received an intrathecal injection containing the equivalent amount of El Kollipher and ethanol, but without paclitaxel.

#### Post-traumatic osteoarthritis (PTOA) via ACL transection

Adult male mice (12 weeks of age) were anesthetized with isoflurane (2–5% in 100% oxygen) as an inhalation anesthetic. Postoperative pain control was provided with buprenorphine (0.05 mg/kg intraperitoneally) on the day of surgery. The left hindlimb area was shaved and disinfected, and sterile surgical drapes were applied. Under magnification with a surgical microscope (Leica M320), the left knee joint was exposed through an anterior skin incision. The joint capsule was opened via medial parapatellar capsulotomy, the patella was dislocated laterally, and with the knee in full flexion, the anterior cruciate ligament (ACL) was identified and transected (ACLx) with microscissors. Following transection, the patella was repositioned and the joint capsule was closed with 7-0 nylon suture. Mechanical allodynia was observed as early as 6 weeks following surgery. Sham-operated control mice underwent identical surgical procedures, except the ACL was not transected. Mice were allowed to recover for 6 days before behavioral testing began, which was performed weekly after ACLx surgery.

### 2.3 Behavioural experiments

Mechanical sensitivity was assessed using the SUDO method with calibrated von Frey filaments (Stoelting) to estimate the 50% withdrawal threshold in force (grams/square millimeter), reported as the paw withdrawal threshold (PWT).[9] Testing was performed in a blinded manner by the same experimenter throughout all cohorts to control for experimenter effects.

For each pain model, 4-9 mice per group were tested. Mice were allowed to habituate for 30 minutes in the testing environment. Baseline mechanical thresholds were established prior to pain induction. Following induction (capsaicin, CFA, SNI, PTOA, or paclitaxel), mechanical thresholds were assessed at appropriate timepoints, as specified in Table 1. Testing areas were chosen to test for secondary hyperalgesia consistent with the pain model as described above, specifically the plantar surface of the hind paw for capsaicin, CFA, and paclitaxel, and the sural edge of the plantar surface of the hind paw for SNI and PTOA. Importantly, we tested away from the original area of insult to assess secondary sensitization. For intrathecal injections, animals that did not produce an obvious tail flick were excluded from analysis as an indication of incomplete injection. Animals were randomly assigned to experimental groups, and the experimenter was blinded during behavioural testing. All mice were tested by the same experimenter and under the same experimental conditions throughout the duration of the experiment. Maximum possible effect (MPE) was used to quantify analgesic effects using the following formula: 100% * [post-treatment paw withdrawal threshold (PWT) – pre-treatment PWT) * (baseline PWT – pre-treatment PWT)^-1^.

**Table 1:** Overview of the pain models used.

|  | **Site and delivery approach for injury/insult** | **Site of Von Frey testing** | **Time of spinal cord tissue collection** |
| --- | --- | --- | --- |
| **Capsaicin** | Intraplantar paw injection, near the walking pads (1% w/v, 5 µl) | Plantar surface, near the heel | 2 hours post injection |
| **Complete Freund's Adjuvant** | Intraplantar paw injection, near the walking pads (100%, 15 µl) | Plantar surface, near the heel | 2 days post injection |
| **Spared Nerve Injury** | Common peroneal and tibial nerve transection | Plantar surface, sural edge | 2 weeks post-surgery |
| **Post-traumatic Osteoarthritis** | Anterior cruciate ligament transection | Plantar surface, sural edge | 19 weeks post-surgery |
| **Paclitaxel** | Intrathecal injection (50 nM, 5 µl) | Plantar surface, near the heel | 2 hours post injection |

### 2.4 Tissue Processing

#### Tissue Extraction

Mice were deeply anesthetized with chloral hydrate (400 mg/kg intraperitoneally) and then transcardially perfused with ice-cold, sucrose-supplemented oxygenated artificial cerebrospinal fluid with kynurenic acid (Sigma-Aldrich, K3375) (sACSF: 51 mM sucrose, 92 mM NaCl, 5 mM KCl, 0.5 mM CaCl₂, 26 mM NaHCO₃, 1.25 mM NaH₂PO₄, 15 mM glucose, 7 mM MgSO₄, 1 mM kynurenic acid on day of experiment; pH 7.4, osmolarity ∼315 mOsm/L). The lumbar spinal cord segment (L4-L6) was rapidly extracted by laminectomy and immediately placed in ice-cold sACSF, then glued against a 4% agar block mounted on a vibratome disk.

The vibratome disk was placed into the vibratome chamber (Leica VT-1200), and 300 μm parasagittal slices were obtained. Slices were then placed in oxygenated sACSF containing OGB-488 AM (Invitrogen, O6807) [22.53 mM DMSO, 0.008% w/v Pluronic F-127] at 34°C for 40 minutes. Osmolarity was maintained at 310-320 mOsm/L. After incubation, slices were removed from solution and transferred to regular ACSF at room temperature for 50 minutes prior to imaging.

In one experiment, slices were incubated in 300 nM Strychnine (Sigma-Aldrich, S0532-25G) and 10 μM Bicuculline (Abcam, ab120110) for one hour to model disinhibition. Strychnine was dissolved in regular ACSF to a stock concentration of 3 μM. Bicuculline was dissolved in regular ACSF to a stock concentration of 100 μM.

#### Imaging Setup

Ca^2+^ imaging was performed on a fixed stage microscope (Olympus, BX5WI) equipped with a water immersion objective at 40x. Excitation light at 488 nm was provided by an LED (ThorLabs, XCITE120). Emitted fluorescence was filtered using a dichroic mirror and captured on an optiMOS CMOS camera (QImaging, Q40862) with a camera adapter (Olympus, U-TV1X-2). Videos were acquired using Ocular (Teledyne, 01-OCULAR-USB) at a frame rate of 1.6 Hz with a spatial resolution of 1920 × 1080 pixels, representing ∼0.32 μm/pixel). Slices were placed in a recording chamber fitted with a harp to stabilize the tissue. Recordings were performed at room temperature, and ACSF was continuously perfused at a rate of 1.0 mL/min.

#### Recording Protocol

After the slice was positioned under the objective, a baseline video was recorded for 2-6 minutes. Glutamate was then added to the perfusate to a final concentration of 10 μM depending on the protocol, and imaging continued for 4 minutes. Subsequently, glutamate concentration was increased to 25 μM and imaging continued for 4 minutes. Each stimulus condition was recorded as a separate video of 6 minutes duration. Slices were allowed to recover between stimulus applications for 10 minutes.

### 2.5 CuMIN Pipeline

#### Code Repository

The repository for this pipeline can be accessed through GitHub using this link: https://github.com/samwcfung/CuMIN.

#### Preprocessing

Regions of interest (ROIs) were manually identified using FIJI (National Institutes of Health, Bethesda, MD, USA) and the ROI Manager tool.[72] ROI polygons were defined to encompass visually apparent neuronal soma and saved in ImageJ .zip format for subsequent automated extraction.

#### Photobleaching and Global Background Correction

Raw image stacks were subjected to photobleaching correction using polynomial detrending. A second-order polynomial was first fitted to the mean fluorescence intensity across all frames to capture broad temporal trends. This was followed by a second-stage correction using a third-order polynomial applied to the residuals. The mean fluorescence value across the entire image stack was calculated and used to normalize corrected frames, preserving relative intensity variations while compensating for exponential decay in signal intensity over time. Condition-specific frame ranges were employed for the detrending procedure (0–500 frames for spontaneous activity conditions; 0–200 frames for stimulus-evoked conditions).

Following photobleaching correction, images were denoised using a median filter with a kernel size of 5×5 pixels. The median filter was applied frame-by-frame to reduce high-frequency noise while preserving cellular boundaries and fine morphological structures.

Background fluorescence was removed using a top-hat morphological transformation with a disk-shaped structuring element of 85-pixel radius. This approach is adapted from MIN1PIPE and Minian, which uses a two-step ‘erosion and dilation’ approach to remove excessively bright background features.[17,59] The top-hat operation was defined as the difference between the original image and its morphological opening, thereby enhancing local bright features while attenuating spatially extended background variations.

#### Motion Correction

Motion correction capabilities were available within the pipeline using the NormCoRRe (Normalization and Motion CORREction) algorithm, which implements both rigid and non-rigid image registration.[27] Rigid motion correction computes whole-frame translations in x and y dimensions for each frame relative to a template image. The template was defined as the mean-intensity projection of the first 30 frames of the preprocessed image stack. For each subsequent frame, a rigid shift was computed by minimizing the squared difference between the frame and the template, with a maximum allowable shift of 10 pixels in either dimension. The computed shift was then applied to translate the entire frame, and the corrected frame was used to update the template via a running average. Non-rigid motion correction addresses localized motion artifacts by dividing the image into patches and computing independent translations for each patch. The image was divided into a grid of overlapping patches, each with dimensions of 50×50 pixels. For each patch and each frame, a local shift was computed relative to the corresponding patch in the template image. Shifts were computed using normalized cross-correlation and were constrained to a maximum displacement of 10 pixels. The corrected image was reconstructed by applying the computed patch-wise shifts and blending overlapping regions to eliminate boundary artifacts.

#### ROI Extraction, Quantification, and Peak-to-Noise Refinement

ROI polygons were automatically extracted from the ImageJ .zip files and rasterized into binary masks matching the dimensions of the preprocessed image stack. Fluorescence time series were extracted by computing the mean pixel intensity within each ROI mask for every frame, yielding raw fluorescence traces F(t) for each ROI.

Global background fluorescence was estimated from pixels exterior to all ROI masks. For each frame, the 20th percentile of background pixel intensities was computed and subtracted from all ROI traces. To correct for slow drift in background signal during the recording, linear regression was performed on the background values within the initial 200 frames, excluding frames with detected activity peaks (prominence >0.05 dF/F). The estimated slope was subtracted from the full background trace, yielding a baseline-corrected background signal that was subsequently subtracted from all ROI traces.

Corrected fluorescence traces were converted to relative change in fluorescence (dF/F) by dividing by a baseline fluorescence value (F₀) calculated from the initial 200 frames. The optimal F₀ value was determined by testing a range of percentiles to identify the one with the most robust and stable baselines across a variety of slices and treatments. For our data, we defined F₀ as the 8th percentile of fluorescence values within this baseline period, thereby providing a robust estimate resistant to transient activity. The resulting dF/F traces were baseline-normalized to zero.

To exclude low-quality ROIs from further analysis, a peak-to-noise ratio (PNR) analysis was performed. For each ROI, the dF/F trace was decomposed into signal and noise components using frequency separation. A Butterworth filter (2nd order) with a cutoff frequency of 0.4 Hz was applied: low-pass filtering yielded the signal component, while high-pass filtering yielded the noise component. The peak-to-noise ratio was defined as the 99th percentile of the signal component divided by the standard deviation of the noise component. ROIs with a PNR value below 8.0 were excluded from subsequent analysis. Signal traces from retained ROIs were smoothed using a moving-average window of 1 frame width.

#### Event Detection and Analysis

Events were detected in individual dF/F traces using a peak-detection algorithm based on the scipy.signal.find_peaks function. Detection parameters were optimized for condition-specific response characteristics:

##### Spontaneous Activity (0 µm stimulus distance)

Peaks were detected across the entire recording period (frames 0–580) using the following parameters: prominence = 0.08 dF/F, peak width = 1 frame, minimum inter-peak distance = 5 frames, minimum height = 0.1 dF/F. ROIs were classified as active if the spontaneous peak frequency exceeded 0.1 events per 100 frames.

##### Stimulus-Evoked Activity (10 µm and 25 µm stimulus distances)

Peaks were detected within the analysis window (frames 230–580, beginning 138 seconds after stimulus onset) using identical detection parameters. ROIs were classified as active if the maximum dF/F value within the analysis window exceeded 0.02 dF/F.

For each detected peak, the following metrics were extracted: peak amplitude, time to peak, rise slope (maximum rate of change during rising phase), decay time (time from peak to 50% decay), peak width at half-maximum prominence, and area under the curve. Rise time was defined as the time required for fluorescence to increase from 10% to 90% of peak amplitude.

ROI traces were subjected to quality control checks to identify motion artifacts and signal degradation. ROIs were flagged if they exhibited: (1) fluorescence variance below 0.01 dF/F^2^, (2) frame-to-frame intensity jumps exceeding 0.5 dF/F, or (3) baseline drift (difference between mean fluorescence in first and last 10% of frames) greater than 30% of the initial mean.

### 2.6 Linear Discriminant Analysis

#### Feature Selection

Using a stepwise approach to determine relative feature weight and importance, a curated set of 18 features was selected to comprehensively characterize neuronal Ca^2+^ dynamics. *Spontaneous Activity Features* (n=3) included spontaneous peak frequency, average peak amplitude, and maximum peak amplitude (spont_peak_frequency_0, spont_avg_peak_amplitude_0, spont_max_peak_0). *Evoked Response Features* were extracted for both 10 μM glutamate (n=4: peak_amplitude_1, peak_area_under_curve_1, peak_max_rise_slope_1, evoked_duration_1) and 25 μM glutamate (n=4: peak_amplitude_2, peak_area_under_curve_2, peak_max_rise_slope_2, evoked_duration_2). *Dose-Response Metrics* (n=7) included delta values (high minus low dose) and ratios (high divided by low dose) for amplitude, area under curve, maximum slope, and duration (Delta_Amp, Delta_AUC, Delta_max_slope, Delta_Duration, Ratio_Amp_10.25, Ratio_AUC, Ratio_max_slope, Ratio_duration).

Peak detection was performed using scipy.signal.find_peaks (SciPy ≥ 1.7.0) with the following parameters: height threshold = 0.1 (normalized dF/F₀), minimum inter-peak distance = 3 frames, minimum prominence = 0.08 dF/F, minimum width = 2 frames, maximum width = 50 frames, and relative height = 0.5 dF/F.

Outliers were identified using the interquartile range (IQR) method applied independently to each feature. Outliers were defined as observations exceeding twice the IQR below the 25th (Q1) or above the 75th (Q3) percentile. Samples exhibiting outlier characteristics in any of the 18 features were excluded from all subsequent analyses. With this criteria, 2 capsaicin, 1 CFA, 5 paclitaxel, and 2 SNI slices were excluded. Following outlier removal, sample sizes were verified to ensure adequate statistical power (minimum n=5 per condition).

#### Feature Standardization

All 18 features were standardized using z-score normalization via scikit-learn’s StandardScaler (≥1.0.0) to ensure equal weighting: z = (x - μ) / σ. Linear discriminant analysis was performed using scikit-learn’s LinearDiscriminantAnalysis class to identify linear combinations of features that optimally separate experimental conditions by maximizing between-class variance relative to within-class variance.

The number of discriminant components was set to min(2, k-1), where k represents the number of classes. LDA was applied only when datasets contained at least two samples per class and at least two distinct classes. Prior probabilities were estimated from class frequencies in the dataset.

#### Model Evaluation and Visualization

Clustering quality was quantified using silhouette scores (range: -1 to +1, higher values indicating better separation) and Davies-Bouldin indices (lower values indicating better clustering), both computed in the LDA-transformed space using scikit-learn functions with Euclidean distance.

LDA projections were visualized as scatter plots color-coded by experimental condition, with 95% confidence ellipses (2 standard deviations) overlaid to illustrate cluster distributions. Ellipse parameters were derived from covariance matrices using eigenvalue decomposition. Explained variance ratios for each discriminant axis were reported to indicate the proportion of between-class variance captured.

All analyses utilized NumPy (≥1.21.0), pandas (≥1.3.0), scikit-learn (≥1.0.0), SciPy (≥1.7.0), matplotlib (≥3.4.0), and seaborn (≥0.11.0).

### 2.7 Immunofluorescence

Immunofluorescence (IF) experiments for PTOA and SNI samples were performed at the University of Calgary. IF experiments for paclitaxel were performed at the University of Toronto.

#### For PTOA and SNI samples

Before tissue collection, mice were euthanized with an overdose of sodium pentobarbital and transcardially perfused with 20 mL of PBS and then cold 4% PFA. Dissected spinal cords were post-fixed in 4% PFA at 4 °C overnight and subsequently allowed to equilibrate in 30% sucrose at 4 °C overnight, before finally embedding in cryoprotectant (VWR Clear Frozen Section Compound) for storage at -80 °C. The lumbar section (L4-L6) of the spinal cord was cut in coronal sections (20 μm) using a cryostat (Leica) and stored at -80 °C until further use.

For immunofluorescent analysis, sections were permeabilized by incubating in 0.5% Triton X-100 in blocking solution (5% bovine serum albumin (BSA) in PBS) for 1 hr at room temperature. Then, sections were treated with primary antibodies (Iba1, rabbit, 019-19741, Wako, 1:200; GFAP, chicken, AB_304558, Abcam, 1:500) in 0.1% Triton X-100 in PBS and left at 4 °C overnight. Subsequently, sections were washed twice in PBS and incubated with secondary antibodies Alexa Fluor 488, 555 or 647 (donkey anti-mouse, rabbit, rat, goat, 1:200, Invitrogen), Hoechst-33258 (1:1000, 14530, Sigma) in PBS for 2h at room temperature, then washed thrice in PBS, stained for nuclei using Hoechst-33258 and mounted using Permafluor mountant. All immunohistochemical stains were confirmed positive with appropriate no primary (secondary alone) controls. Image collection and quantification was done using either Nikon A1 or a Leica SP8 confocal microscope. Montages were collected using a 20x objective lens, Tile-Scan and Z stack (8 z-planes) features, and maximum projection images. All imaging of a given stain was performed using the same laser settings. We used ImageJ and designated markers to quantify fluorescence intensity and count cells or cellular components of interest.

#### For paclitaxel samples

Before tissue collection, mice were deeply anesthetized with chloral hydrate (400 mg/kg intraperitoneally) and then transcardially perfused with ice-cold, sucrose-supplemented oxygenated artificial cerebrospinal fluid (sACSF) with kynurenic acid (Sigma-Aldrich, K3375) (sACSF: 51 mM sucrose, 92 mM NaCl, 5 mM KCl, 0.5 mM CaCl₂, 26 mM NaHCO₃, 1.25 mM NaH₂PO₄, 15 mM glucose, 7 mM MgSO₄]; pH 7.4, osmolarity ∼315 mOsm/L). The lumbar spinal cord segment (L4-L6) was rapidly extracted by laminectomy and immediately placed post-fixed in 4% PFA at room temperature for 2 hours, before being washed in 1x PBS and equilibrated in 30% sucrose (w/v) at 4% until it sank to the bottom. Samples were embedded in cryoprotectant (VWR Clear Frozen Section Compound) and sectioned using a cryostat (Leica) into coronal sections (20 μm) and stored at -20 °C until further use.

For immunofluorescent analysis, sections were permeabilized by washing 5 x 5 minutes in 0.1% PBS-T (1 x PBS, 0.1% Triton-X) at room temperature. Sections were then blocked in 10% horse serum (Life Technologies, 16050122) in PBS-T for 1 hour at room temperature, before incubating in primary antibodies diluted in blocking solution (IBA1, mouse, 234001, Synaptic Systems, 1:1000; GFAP, rabbit, ab7260, Abcam, 1:1000) overnight at room temperature. The next day, sections were washed 5 x 5 minutes in PBS-T and incubated in secondary antibody (Alexa Fluor 488, 567 or 647 (donkey anti-mouse or rabbit, 1:500, Jackson Immunoresearch) for 2 hours at room temperature in the dark, followed by 5 x 5 minutes washes in PBS-T. DAPI (1:500 in PBS) was applied for 10 minutes at room temperature and immediately washed off with 2 x 5 minutes with PBS, then mounted with Daco or Prolong Gold Antifade (ThermoFisher, P36930). Slides were sealed with clear nail polish and stored in 4 °C until imaging. Image collection was done using a Zeiss confocal microscope (Zeiss, Observer.Z1 with LSM 700) using a 20x objective lens with Z-stacks performed at every 2-3 µm, combined with tile scans to capture each dorsal horn unilaterally. All imaging of a given stain was performed with the same laser settings. FIJI was used to quantify fluorescence intensity, and Sholl analysis was performed with the “Sholl Analysis” plugin developed by Ferreira et al.[23]

### 2.8 Statistical Analysis

All statistical analyses were performed using GraphPad Prism (v10) and Python (v3.11). Data are presented as the mean ± standard error of the mean (SEM). Fisher’s exact tests, odds ratios (OR), one-and two-way ANOVAs with post-hoc testing, one- and two-sided student’s t-tests, Mann-Whitney U tests, and Pearson’s R correlation testing were used as appropriate, with α set to 0.05. A priori sample sizes were determined with previous preliminary data from our lab.

## Results

### 3.1 Development of a Ca^2+^ imaging and analysis pipeline to investigate central changes in mouse pain models

To investigate central changes in spinal cord cellular activity during pathological pain, we began by developing a method for Ca^2+^ imaging of spinal cord neurons using epifluorescent imaging and *ex-vivo* spinal cord slices (Figure 1A). After a rapid dorsal laminectomy and incubation of lumbar spinal cord slices in OG 488 BAPTA-1 AM, we imaged the tissue using an epifluorescent microscope to observe changes in intracellular Ca^2+^ dynamics. While we were able to quantify changes in fluorescence manually using Clampfit, this method was time consuming and required user-defined cutoffs and background subtraction that can introduce bias.

**Figure 1.**
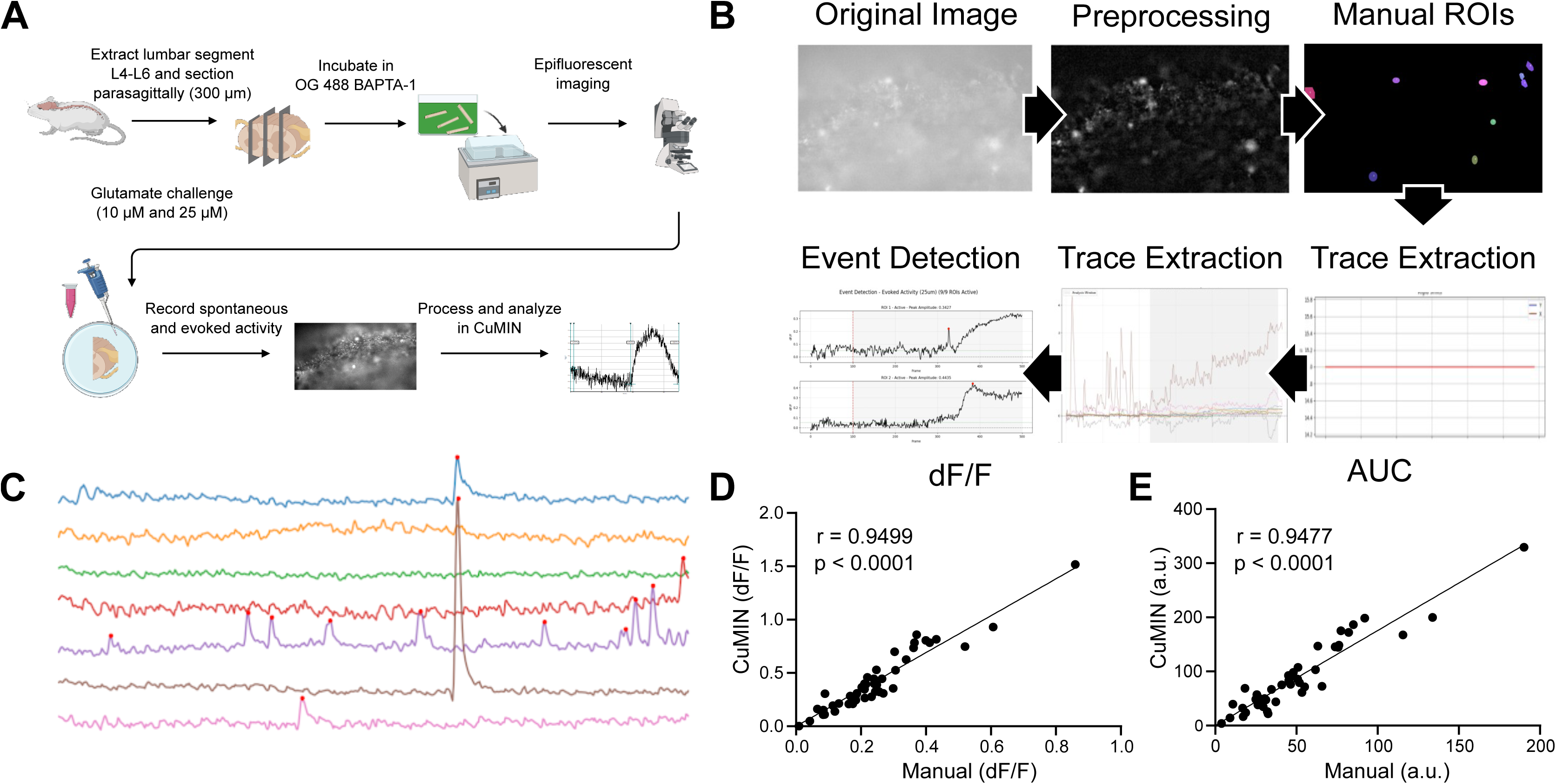
CuMIN: A pipeline to analyze Ca^2+^ imaging data. **A** Schematic of Ca^2+^ imaging and analysis workflow, including dissection, Ca^2+^-sensitive dye (OG 488 BAPTA-1 AM), epifluorescent imaging, glutamate challenge, and processing in CuMIN. Created in BioRender: https://BioRender.com/vhhmh2k **B** Overview of the CuMIN image processing pipeline, encompassing preprocessing, manual ROI selection, motion correction, trace extraction, and event detection. **C** Sample spontaneous activity traces from a model of disinhibition: strychnine (300 nM) and bicuculline (10 μM). Each colour represents a separate ROI. **D** Validation of dF/F from CuMIN against manual analysis. n=45 cells/3 animals. Statistics: Pearson’s r. r=0.95, p < 0.0001. **E** Validation of AUC from CuMIN against manual analysis. n=45 cells/3 animals. Statistics: Pearson’s r. r=0.95, p < 0.0001.

We next turned to publicly available analysis pipelines, such as MIN1PIPE, Minian, and CaImAn, which were developed for 1-photon miniscope applications and analysis.[17,27,59] These pipelines have been used extensively for miniscope applications, but they lack optimization for epifluorescent applications and manual ROI curation strategies in approaches where image quality is a concern. Specifically, we experienced x-, y-, and z-drift over the imaging duration, causing distortion to regions of interest (ROIs). As such, ROIs that were detected using principal component analysis (PCA) or constrained non-negative matrix factorization (CNMF) approaches commonly used in established pipelines did not accurately capture the region of interest.[27,65,69] Furthermore, cross-session registration was compromised as positional drift caused ROIs to shift.

To overcome this limitation, we employed the gold standard approach of manually defining ROIs and built a custom pipeline using Python based on two previous pipelines, MIN1PIPE and Minian.[17,59] Using elements of background subtraction, motion correction, and signal refinement for these pipelines, and adding a custom built module to integrate manually selected ROIs, we created our own pipeline called CuMIN (**Cu**rated ROIs for **MIN**ian). A brief workflow is depicted in Figure 1B, showing sample images of image preprocessing, which includes background subtraction and photobleaching correction, followed by motion correction, then trace extraction of manually selected ROIs, and finally event detection. This pipeline was created with the intention to be as modular as possible for the end user, allowing for parameters to be adjustable and each visualized in real time using Jupyter Notebook, enabling rapid optimization for tissue-specific and condition-specific recordings (Figure 1C-E).

### 3.2 Validating CuMIN using a model of disinhibition

We validated CuMIN through technical and biological approaches. We performed technical validation by comparing manually-extracted Ca^2+^ traces to pipeline outputs and found a high degree of correlation (r > 0.94) for dF/F and AUC calculations when comparing manually analyzed traces in Excel and Clampfit against our CuMIN pipeline (Figure 1C-E). We followed this up with biological validation using a model of disinhibition through inhibition of GABAergic and glycinergic signalling (strychnine and bicuculline).

We first observed that some cells exhibited spontaneous Ca^2+^ events during naive conditions, which we termed spontaneously active (SA). Another population of cells were not spontaneously active but would exhibit a Ca^2+^ response when exposed to the excitatory neurotransmitter glutamate via the perfusate, which we termed non-spontaneously active (NSA). We decided to separately analyze SA from NSA cells to determine if there were changes in spontaneous and evoked behaviours in these two populations. We next predicted that there would be increased glutamate-evoked recruitment of non-spontaneously active (NSA) cells, as previous studies have established that excitatory SDH neurons are especially disinhibited by chloride dysregulation.[49,83] We found that spontaneous network activity was unchanged in disinhibited tissue when comparing the percentages of SA with NSA cells (Figure 2A). However, we observed a significant increase in the number of NSA cells that responded when exposed to the excitatory neurotransmitter glutamate (Figure 2B). We observed no changes in the frequency and amplitude of spontaneous events (Figure 2C-D). These findings suggest that there is a facilitated increase in the responsiveness of NSA cells, potentially along with the recruitment of a previously silent population to the NSA network. These findings also indicate that altered NSA recruitment during disinhibition could be observed in our imaging and analysis approach.

**Figure 2.**
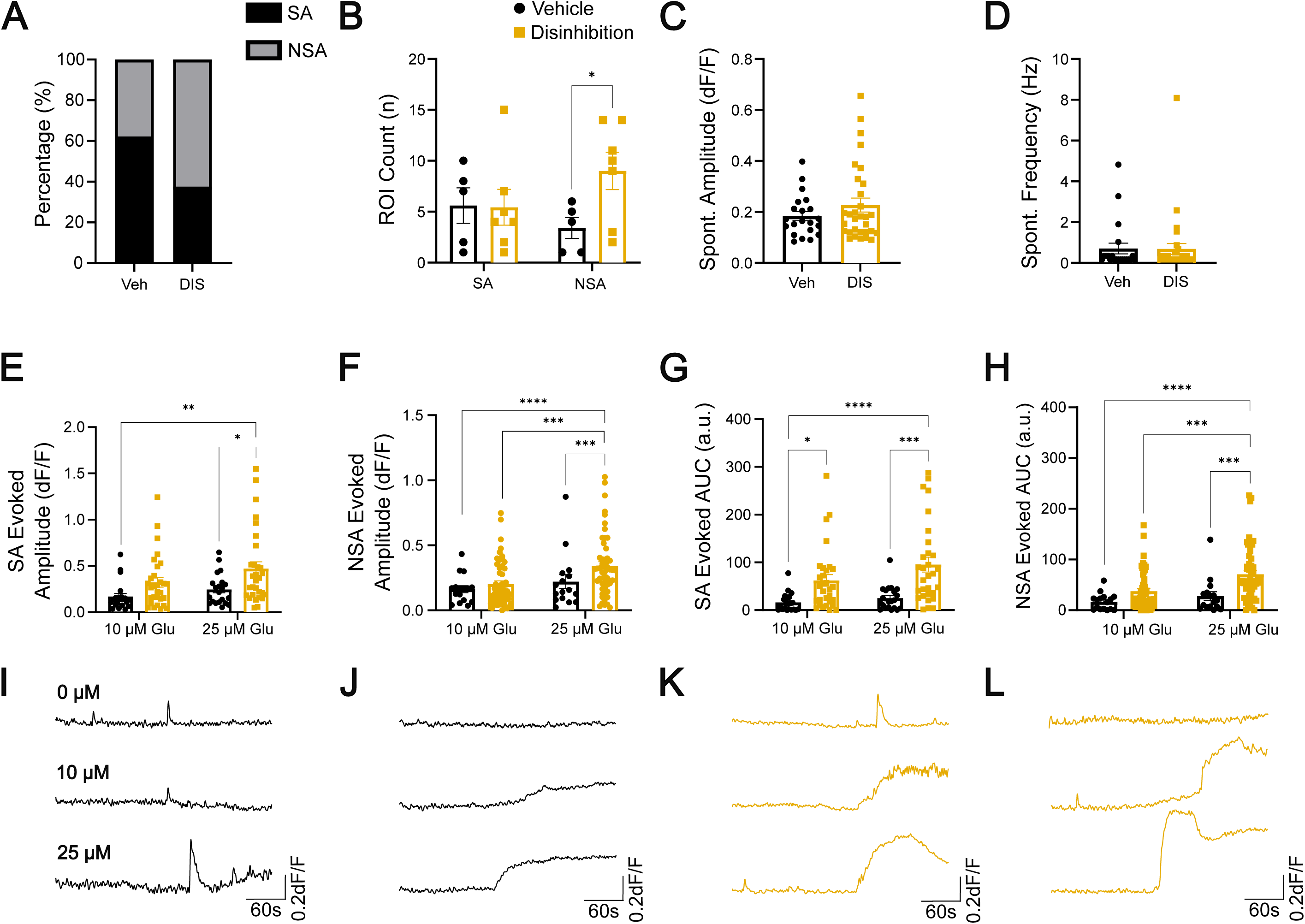
Validation of CuMIN using a model of disinhibition induced by strychnine (300 nM) & bicuculline (10 μM). **A** Total percentages of cells that exhibit spontaneous activity (in black, SA) compared to cells that only respond to glutamate (in grey, NSA) out of all cells that respond to glutamate. n=37-83 cells/3-5 animals/group. Statistics: Fisher’s exact test, OR=2.202, p=0.072. **B** Comparison of the number of SA and NSA cells between vehicle and disinhibition treated tissue. n=5-7 slices/3-5 animals/group. Statistics: 2-way ANOVA with Fisher’s LSD. Cell activity effect: F(1,20)=0.16, p=0.70; treatment effect: F(1,20)=0.24, p=0.13; interaction effect: F(1,20)=2.76, p=0.11. **C** Average amplitude of spontaneous events from spontaneously active cells. n=21-31 cells/3-4 animals/group. Statistics: Mann-Whitney U test, U=298, p=0.62. **D** Average frequency of spontaneous events from spontaneously active cells. n=21-31 cells/3-4 animals/group. Statistics: Mann-Whitney U test, U=324.5, p=0.99. **E** The amplitude of response from spontaneously active (SA) cells under glutamate challenge at 10 μM and 25 μM. n=16-52 cells/3-4 animals/group. Statistics: 2-way ANOVA with Tukey’s post-hoc. Glutamate effect: F(1,100)=3.67, p=0.058; treatment effect: F(1,100)=8.91, p=0.0036; interaction effect: F(1,100)=0.96, p=0.32. **F** The amplitude of response from non-spontaneously active (NSA) cells under glutamate challenge at 10 μM and 25 μM. n=16-52 cells/3-4 animals/group. Statistics: 2-way ANOVA with Tukey’s post-hoc. Glutamate effect: F(1,132)=6.001, p=0.016; treatment effect: F(1,132)=3.98, p=0.048; interaction effect: F(1,132)=0.98, p=0.32. **G** The area under the curve from SA cells under glutamate challenge at 10 μM and 25 μM. n=16-52 cells/3-4 animals/group. Statistics: 2-way ANOVA with Tukey’s post-hoc. Glutamate effect: F(1,98)=3.031, p=0.085; treatment effect: F(1,98)=23.32, p<0.0001; interaction effect: F(1,98)=0.99, p=0.32. **H** The area under the curve from NSA cells under glutamate challenge at 10 μM and 25 μM. n=16-52 cells/3-4 animals/group. Statistics: 2-way ANOVA with Tukey’s post-hoc. Glutamate effect: F(1,132)=6.46, p=0.012; treatment effect: F(1,132)=13.33, p=0.0004; interaction effect: F(1,312)=1.62, p=0.20. **I** Representative traces from a SA cell pre-treated with vehicle and exposed to 0, 10, and 25 μM glutamate. **J** Representative traces from a NSA cell pre-treated with vehicle and exposed to 0, 10, and 25 μM glutamate. **K** Representative traces from a SA cell pre-treated with strychnine and bicuculline and exposed to 0, 10, and 25 μM glutamate. **L** Representative traces from a NSA cell pre-treated with strychnine and bicuculline and exposed to 0, 10, and 25 μM glutamate. **B-L** Vehicle control (ACSF) shown in black, disinhibition shown in gold. For all statistics, *p<0.05, **p<0.01, ***p<0.001, ****p<0.0001. Error bars represent SEM.

To mimic hyperalgesia, which is behaviour that occurs when a normally painful stimulus evokes an exaggerated pain response in vivo, we bath-applied 10 µM and 25 µM glutamate to mimic biologically relevant concentrations of extracellular glutamate during these types of stimuli ex vivo.[54,56,97] Using this glutamate-evoked approach, we further observed that disinhibited cells have a higher evoked response amplitude (dF/F) and area under the curve (AUC) compared to vehicle (VEH) controls (Figure 2E-L). This effect was observed in both SA and NSA cells when exposed to 25 μM glutamate. Together, these data suggest that during disinhibition there is: (a) a recruitment of a previously non-responsive population to the NSA network, and (b) an increase in the magnitude of glutamate evoked Ca^2+^ responses in both SA and NSA cells. In summary, these findings indicate that our model of disinhibition replicated similar findings from *in-vivo* models of neuropathic pain, in which glutamate-evoked Ca^2+^ responses were increased in amplitude,[83] as well as in *ex-vivo* models of disinhibition, where cells exhibit an enhanced response to stimulation, confirming that our pipeline faithfully captures previously characterized physiology.[18]

### 3.3 Spontaneous activity in spinal nociceptive networks is altered in CFA and SNI models of pathological pain

We next examined if acute and chronic pathological pain models exhibited unique patterns of spontaneous Ca^2+^ events in cells by testing tissue after insult by capsaicin (CAP), complete Freund’s adjuvant (CFA), spared nerve injury (SNI), post-traumatic osteoarthritis (PTOA), and paclitaxel (PAC) to observe changes under baseline conditions (Figure 3). While we did not observe changes in cellular activity following Capsaicin, an acute model of inflammatory pain (Figure 3Ai-Av), we observed several changes in activity following CFA, a more protracted model of inflammatory pain (Figure 3Bi-Bv). Notable, we observed a significant increase in the proportion of NSA cells following CFA compared to their contralateral controls (Figure 3Bi). This effect was driven by an overall increase in the number of NSA cells, with no changes to the number of SA cells (Figure Bii). The opposite effect was observed in the proportion of NSA cells following SNI, where we observed an increase in the number of SA cells (Figure 3Ci). No significant changes in relative proportions, population counts, or spontaneous activity dynamics were observed for PTOA (Figure 3Di-Dv) or paclitaxel (Figure 3Ei-Ev). We did note a trend of more cells recruited to both the SA and NSA populations for paclitaxel, but this was nonsignificant (p = 0.11) (Figure 3Eii). These findings suggest differential recruitment during CFA and SNI, where the NSA network recruits additional cells in CFA; while during SNI, there is a shift towards SA cells. These divergent changes in network activity between the CFA and SNI models are unsurprising, given their vastly different etiology.[14,24,26,39,44,47]

**Figure 3.**
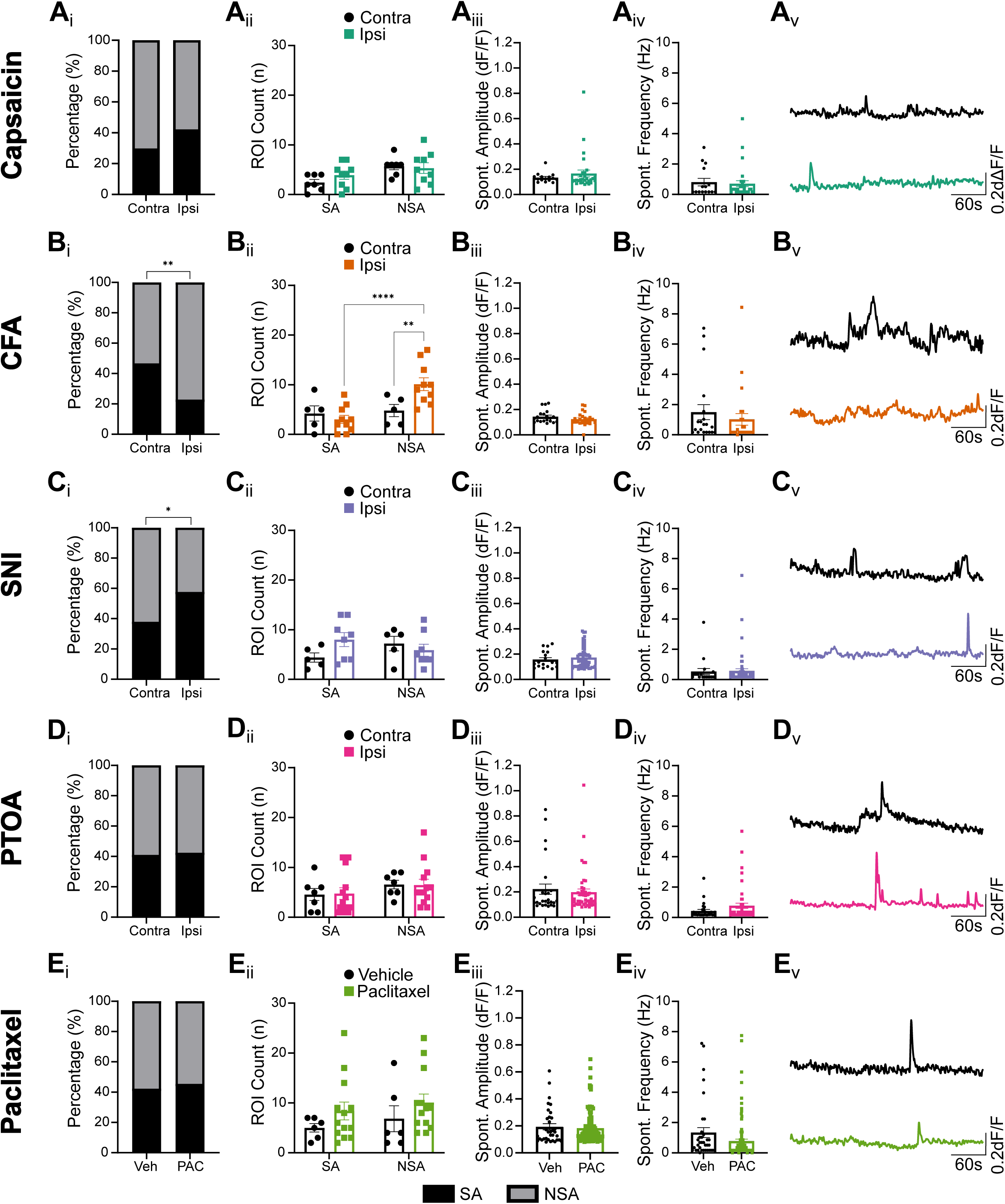
Spontaneous activity parameters for acute and chronic pathological pain models, Capsaicin, CFA, SNI, PTOA, and Paclitaxel. Ai-Ei: Total percentages of cells that are Spontaneously Active (SA) compared to cells that are Non-spontaneously Active (NSA) and only fluoresce in the presence of glutamate are quantified. n=15-184 cells/4-8 animals/group. Statistics: Fisher’s exact test. **Ai:** OR=0.56, p=0.18; **Bi:** OR=3.45, p=0.0022; **Ci:** OR=0.43, p=0.018; **Di:** OR=0.94, p=0.88; **Ei:** OR=0.97, p>0.99. **Aii-Eii:** A comparison of cellular activity of SA and NSA cells separated by treatment and level of activity. n=7-13 slices/4-8 animals/group. Statistics: 2-way ANOVA with Fisher’s LSD. **Aii** Cell activity effect: F(1,28)=7.34, p=0.011; treatment effect: F(1,28)=0.38, p=0.54; interaction effect: F(1,28)=1.11, p=0.30. **Bii** Cell activity effect: F(1,26)=8.90, p=0.0061; treatment effect: F(1,26)=2.52, p=0.12; interaction effect: F(1,26)=6.35, p=0.018. **Cii** Cell activity effect: F(1,22)=0.061, p=0.81; treatment effect: F(1,22)=0.69, p=0.41; interaction effect: F(1,22)=3.25, p=0.085. **Dii** Cell activity effect: F(1,36)=2.14, p=0.15; treatment effect: F(1,36)=0.0012, p=0.97; interaction effect: F(1,36)=0.015, p=0.90. **Eii** Cell activity effect: F(1,34)=0.74; treatment effect: F(1,34)=2.60; interaction effect: F(1,34)=0.0012, p=0.97. **Aiii-Eiii:** The average amplitude of spontaneous events from spontaneously active cells. n=15-105 cells/4-8 animals/group. Statistics: Mann-Whitney U test. **Aiii** U=187, p=0.46; **Biii** U=179, p=0.16; **Ciii** U=476, p=0.65; **Diii** U=597, p=0.79; **Eiii** U=1755, p=0.82. **Aiv-Eiv:** The average frequency of spontaneous events from spontaneously active cells. n=15-105 cells/4-8 animals/group. Statistics: Mann-Whitney U test. **Aiv** U=216, p=0.83; **Biv** U=186.5, p=0.19; **Civ** U=485.5, p=0.72; **Div** U=554, p=0.43; **Eiv** U=1478, p=0.075. **Av-Ev:** Representative spontaneous activity traces for each pain model. For all statistics, *p<0.05, **p<0.01, ***p<0.001, ****p<0.0001.

### 3.4 Glutamate-evoked activity is altered in capsaicin and PTOA models of pathological pain

We next asked if glutamate-evoked activity was altered in SA and NSA populations after exposure to 10 µM and 25 µM glutamate (Figure 4). These concentrations were chosen to mimic physiologically relevant extracellular concentrations during stimuli. Starting with SA cells, we again found no significant alterations in SA cells following capsaicin, a model of acute inflammation (Figure 4Ai-Aiii). However, we observed that NSA cells exposed to capsaicin tended to have a higher, but non-significant, amplitude of response at 10 µM glutamate, compared to their contralateral controls (Figure 4Aiv). When we compared the ratio of the response at 10 µM to 25 µM glutamate for these NSA cells, we discovered that ipsilateral cells had a higher ratio of response compared to contralateral cells (Figure 4Av). This indicated that the responses of ipsilateral NSA cells were achieving a ceiling effect in their amplitude of response at a lower dose of glutamate, suggesting an increase in glutamate-evoked responsiveness. In our more protracted form of inflammatory pain, CFA, we observed no significant changes in either SA or NSA populations following exposure to glutamate, suggesting that the central changes following CFA are primarily driven by changes in spontaneous, not evoked activity. Following SNI, we observed a reduced amplitude for ipsilateral SA cells (Figure 4Ci), with no changes to NSA cells (Figure 4Civ-Cvi), indicating a reduction in evoked Ca^2+^ signaling in SA cells. Conversely, we observed an increased amplitude for ipsilateral SA cells following PTOA (Figure 4Di), along with an increased ratio of response at 10 µM to 25 µM glutamate, indicating that ipsilateral NSA cells were achieving a ceiling effect in response to glutamate (Figure 4Div-Dvi). Similar to what we observed for SNI, we observed a reduced amplitude for ipsilateral cells from cells treated by paclitaxel (Figure 4Ei), indicating a reduction in evoked Ca^2+^ signaling. However, we also observed an increase in the amplitude of NSA cells treated by paclitaxel in response to glutamate, compared to their ipsilateral controls (Figure 4Eiv). Altogether, we observed that glutamate-evoked activity in SA populations from capsaicin and CFA, as well as NSA populations from CFA and SNI are relatively robust and stable following insult. We found that glutamate-evoked activity in NSA populations is more likely to be perturbed, such that activity profiles in SA populations from SNI, PTOA, and paclitaxel, as well as NSA populations from capsaicin, PTOA, and paclitaxel exhibit altered responsiveness following insult or injury.

**Figure 4.**
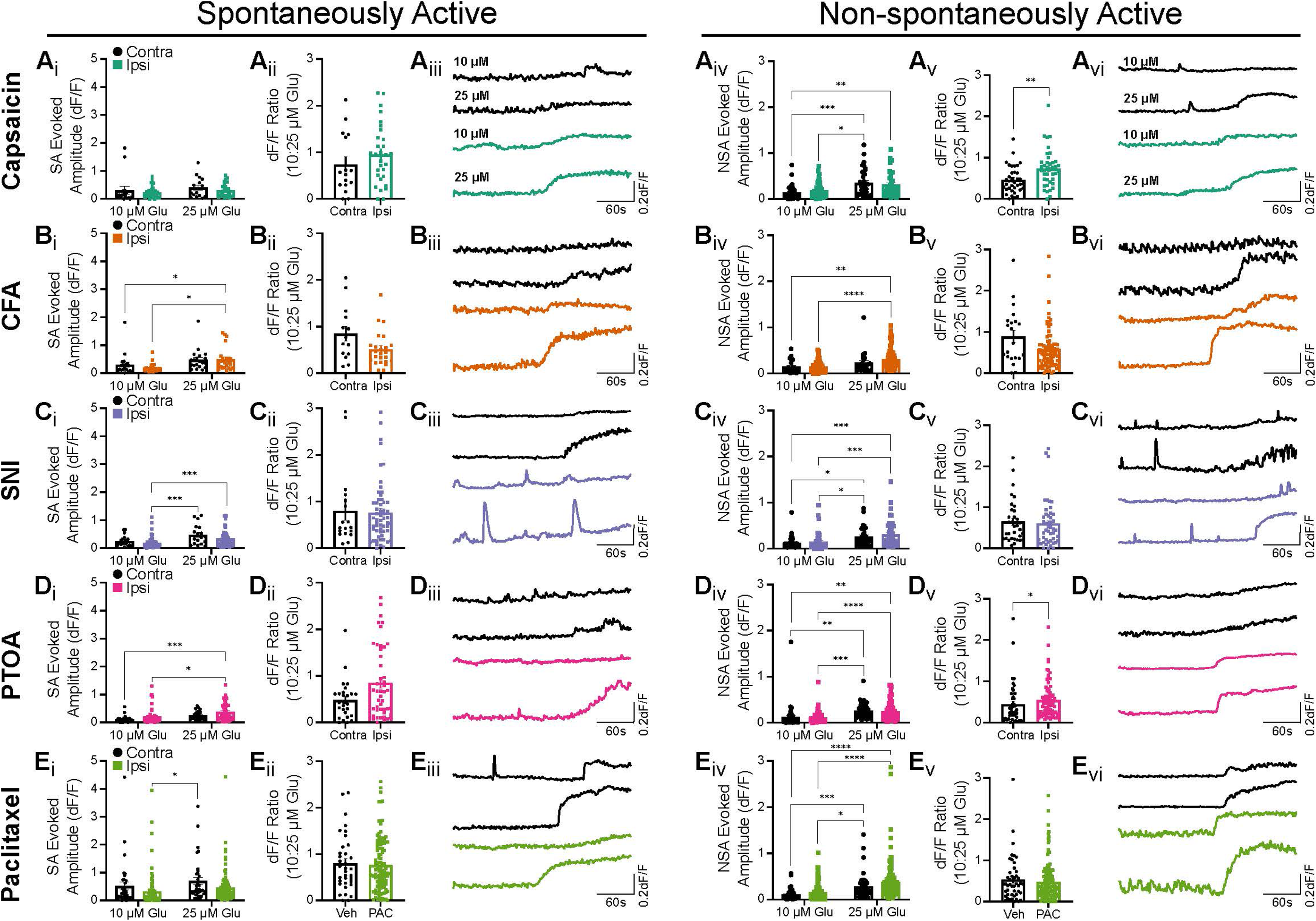
Glutamate evoked activity parameters for acute and chronic pathological pain models, Capsaicin, CFA, SNI, PTOA, and Paclitaxel. Ai-Ei: The amplitude of response from spontaneously active (SA) cells under glutamate application (10 μM and 25 μM). n=15-109 cells/4-8 animals/group. Statistics: 2-way ANOVA with Tukey’s post-hoc. **Ai** Glutamate effect: F(1,88)=1.38, p=0.24; treatment effect: F(1,88)=1.49, p=0.22; interaction effect: F(1,88)=0.033, p=0.86. **Bi** Glutamate effect: F(1,82)=9.98, p=0.0022; treatment effect: F(1,82)=0.42, p=0.52; interaction effect: F(1,82)=0.78, p=0.38. **Ci** Glutamate effect: F(1,146)=13.45, p=0.0003; treatment effect: F(1,146)=3.97, p=0.048; interaction effect: F(1,146)=0.60, p=0.44. **Di** Glutamate effect: F(1,142)=11.44, p=0.0009; treatment effect: F(1,142)=7.062, p=0.0088; interaction effect: F(1,142)=0.00012, p=0.99. **Ei** Glutamate effect: F(1,276)=3.39, p=0.067; treatment effect: F(1,276)=6.55, p=0.011; interaction effect: F(1,276)=0.044, p=0.83. **Aii-Eii:** For SA cells, the ratio of the amplitude of response at 10 μM glutamate to the response at 25 μM. n=15-109 cells/4-8 animals/group. Statistics: Mann-Whitney U. **Aii** U=188, p=0.23; **Bii** U=107, p=0.067; **Cii** U=480, p=0.69; **Dii** U=462, p=0.070; **Eii** U=1797, p=0.78. **Aiii-Eiii:** Representative traces from ipsilateral and contralateral SA cells under glutamate application. **Aiv-Eiv:** The amplitude of response from non-spontaneously active (NSA) cells under glutamate application (10 μM and 25 μM). n=20-134 cells/4-8 animals/group. Statistics: 2-way ANOVA with Tukey’s post-hoc. **Aiv** Glutamate effect: F(1,148)=19.59, p<0.0001, treatment effect: F(1,148)=0.030, p=0.86; interaction effect: F(1,148)=1.36, p=0.24. **Biv** Glutamate effect: F(1,212)=16.26, p<0.0001; treatment effect: F(1,212)=1.811, p=0.18; interaction effect: F(1,212)=1.72, p=0.19. **Civ** Glutamate effect: F(1,152)=16.22, p<0.0001; treatment effect: F(1,152)=0.99, p=0.32; interaction effect: F(1,152)=0.14, p=0.71. **Div** Glutamate effect: F(1,230)=30.72, p<0.0001; treatment effect: F(1,230)=0.30, p=0.58, interaction effect: F(1,230)=0.0011, p=0.97. **Eiv** Glutamate effect: F(1,358)=32.66, p<0.0001; treatment effect: F(1,358)=4.48, p=0.035; interaction effect: F(1,358)=0.28, p=0.60. **Av-Ev:** For NSA cells, the ratio of the amplitude of response at 10 μM glutamate to the response at 25 μM. n=20-134 cells/4-8 animals/group. Statistics: Mann-Whitney U. **Av** U=451.5, p=0.0048; **Bv** U=680, p=0.070; **Cv** U=673, p=0.49; **Dv** U=1205, p=0.035; **Ev** U=2960, p=0.64. **Avi-Evi:** Representative traces from ipsilateral and contralateral SA cells under glutamate application. For all statistics, *p<0.05, **p<0.01, ***p<0.001, ****p<0.0001. Error bars represent SEM.

### 3.5 Supervised clustering reveals unique Ca^2+^ signatures in models of pathological pain

Having established CuMIN’s capacity to extract diverse features from Ca^2+^ imaging data, we were left with a fundamental question: how do these features collectively distinguish one pain model from another? The sheer breadth of our dataset, spanning temporal, spatial, and network-level descriptors, demanded a dimensionality reduction strategy that could reveal underlying patterns without sacrificing biological interpretability. Specifically, we wondered whether models with shared underlying mechanisms would cluster together in feature space, and whether such clustering could provide insight into less characterized conditions like PTOA. To do so, we extracted a number of additional features, including a total of 18 activity, response, and dose-response features, to perform dimension reduction using a linear discriminant analysis (LDA) (Figure 5). We reasoned that CFA, capsaicin (CAP), paclitaxel (PAC), and SNI were well established pain models in the field, with known central mechanisms. However, our model of ACL transection, PTOA, has not been extensively studied, such that its central changes are not well understood. As such, we decided to split our data into a train-test approach, training our LDA to cluster CFA, CAP, PAC, and SNI, and then testing our model to superimpose the data points for PTOA onto the LDA space. We first observed clustering of CAP and CFA data points, suggesting similar central mechanisms were present for both inflammatory models. SNI was clustered distinctly along LD1, which is unsurprising given it is a neuropathic pain model involving disinhibition and microglial activation.[14,15,63] Interestingly, PAC had some overlap with CFA, with both models having documented SDH astrocytic activation being a shared mechanistic hallmark.[24,26,41,51,57,68,86,92,100] When we tested our model with PTOA, we observed a significantly more widespread distribution, with significant overlap visible with PAC, and to a lesser extent, SNI. These findings suggested that the central mechanisms present in PTOA might contain elements from both PAC and SNI.

**Figure 5.**
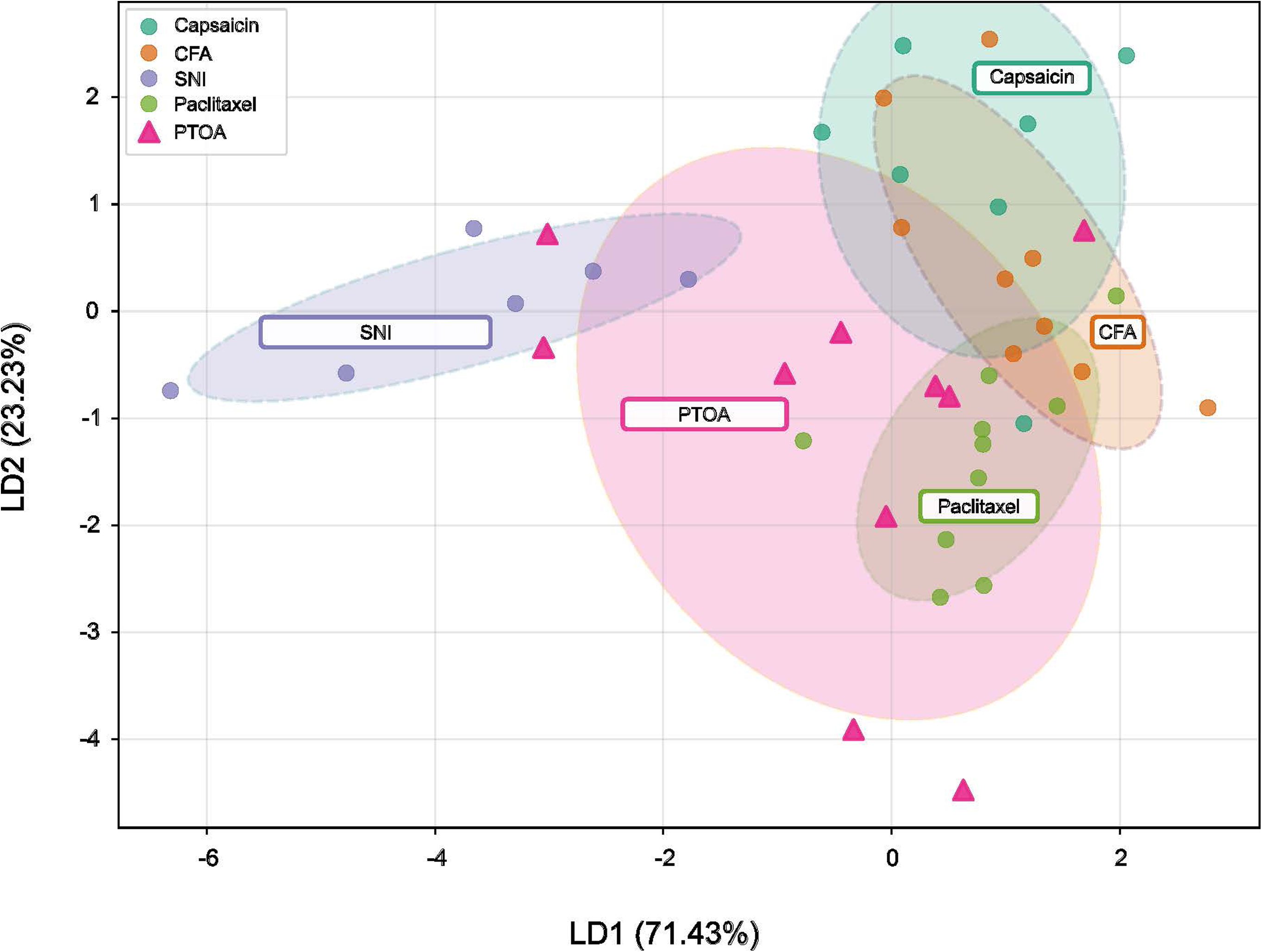
Supervised clustering reveals distinct Ca^2+^ signatures. Supervised clustering using an LDA train-test approach indicates clustering of nociceptive pain models (capsaicin and CFA) together, with neuropathic pain (SNI) separated along LD1 (71.43% variance). Paclitaxel clusters on LD1 along with capsaicin and CFA with some separation along LD2 (23.23% variance). PTOA overlaps with the SNI and paclitaxel clusters. n=6-10 slices/4-8 animals/group.

### 3.6 Astro- and microgliosis is present in PTOA

Based on our findings using LDA to analyze our Ca^2+^ imaging data, we next hypothesized that targeting shared elements of PAC and SNI in the PTOA model might be analgesic. Glial activation, particularly astrocytes and microglia, have been firmly established as key players involved in the maintenance of mechanical allodynia in PAC and SNI. Previous studies have established that SDH astrogliosis is present after systemic and intrathecal administration of PAC, and suppressing this activation can be analgesic.[51,57,98] Conversely, SDH microgliosis has been firmly established in SNI and other neuropathic pain models, and abrogating microglial activity using pharmacological and genetic approaches has been shown to attenuate hypersensitivity.[14,63,85] Peripheral inflammation has been documented to contribute to the maintenance of mechanical hypersensitivity in PTOA, with some evidence of glial involvement.[28,61,62,74]

We therefore performed immunofluorescence to assess the degree of astrocytic and microglial activation in our PAC and PTOA models (Figure 6). In our PAC model, we observed an upregulation of GFAP, indicative of astrocytic activation (Figure 6A,C). We next quantified microglial activation using a marker of microglia, IBA1. We found no observable increase in total IBA1 intensity in the SDH for our PAC model in intensity (Figure 6AB,C). However, previous studies have suggested that microglial activation can come in stages that are not easily quantified by intensity alone, such as increased ramification in a heightened surveillance state.[67] As such, we next quantified the degree of microglial branching using Sholl analysis.[5] Similar to previous studies, we did not observe an increase in the number of radial or total intersections in the SDH following PAC (Figure 6D-F).[51,57,98]

**Figure 6.**
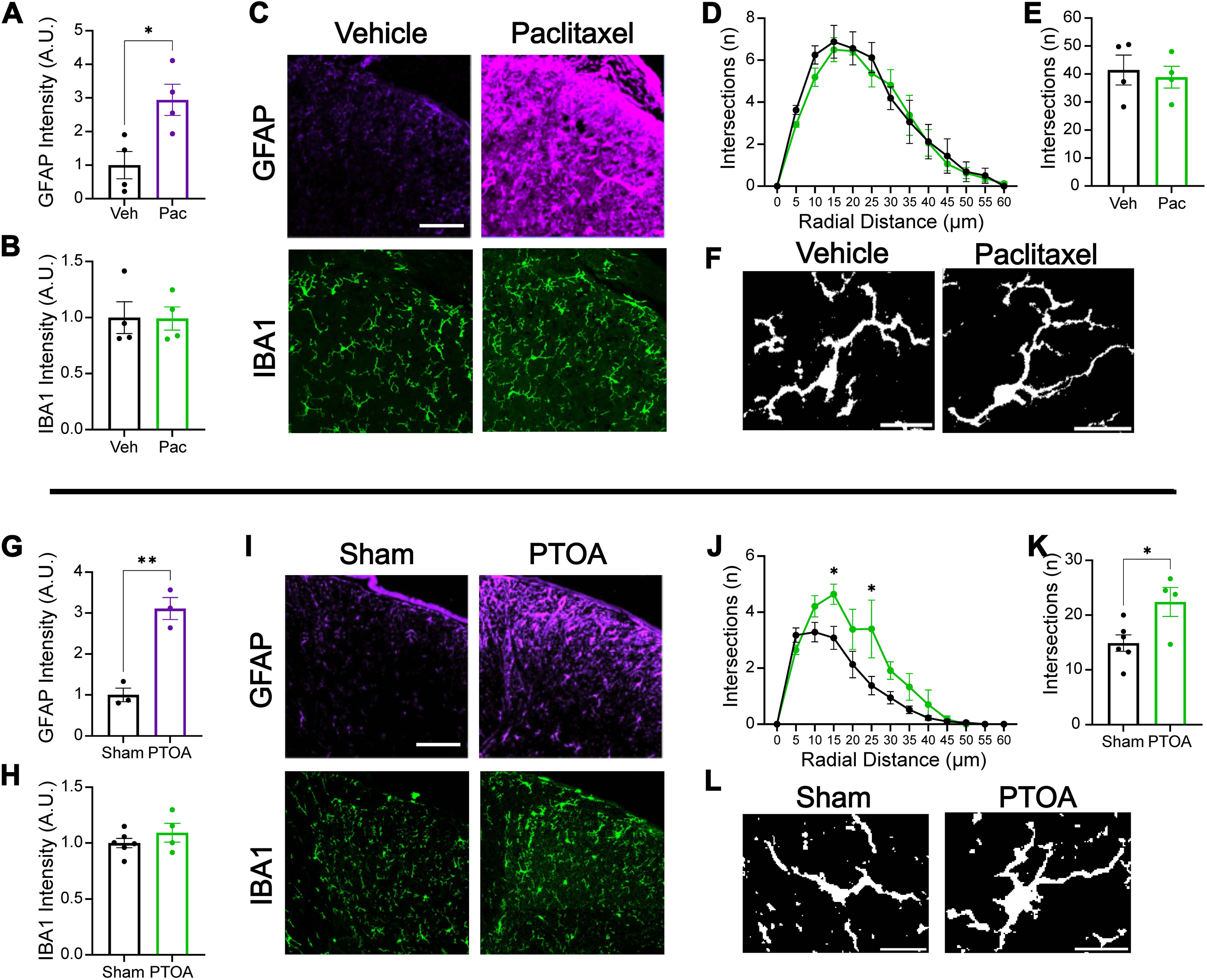
Characterization of gliosis in paclitaxel and PTOA. **A-F:** The effects of intrathecal paclitaxel (50 nM, 4 μL) on gliosis in the dorsal horn. **A:** Normalized mean GFAP intensity to quantify astrogliosis. n=4 animals/group. Statistics: Unpaired t-test. t=3.153, p=0.020. **B:** Normalized mean IBA1 intensity to quantify microgliosis. n=4 animals/group. Statistics: Unpaired t-test. t=0.047, p = 0.96. **C:** Representative images of GFAP (purple) and IBA1 (green). Scale bar indicates 50 μm. **D:** Sholl analysis of microglia, counting intersections at 5 μm stepwise circles to quantify neurite spread. n=3-6 cells/4 animals/group. Statistics: 2-way ANOVA with Sidak’s post-hoc. Distance effect: F(12,72)=65.64, p<0.0001; treatment effect: F(1,6)=0.15, p=0.71; interaction effect: F(12,72)=0.53, p=0.89. **E:** Total average intersections for microglia. n=3-6 cells/4 animals/group. Statistics: Unpaired t-test. t=0.39 p=0.71. **F:** Representative microglial images. Scale bar indicates 10 μm. **G-L:** The effects of PTOA on gliosis in the dorsal horn. **G:** Normalized mean GFAP intensity to quantify astrogliosis. n=4-6 animals/group. Statistics: Unpaired t-test. t=6.71, p=0.0026. **H:** Normalized mean IBA1 intensity to quantify microgliosis. n=4-6 animals/group. Statistics: Unpaired t-test. t=1.092, p=0.31. **I:** Representative images of GFAP (purple) and IBA1 (green). Scale bar indicates 50 μm. **J:** Sholl analysis of microglia, counting intersections at 5 μm stepwise circles to quantify neurite spread. n=3-6 cells/4-6 animals/group. Statistics: 2-way ANOVA with Sidak’s post-hoc. Distance effect: F(12, 96)=50.55, p<0.0001; treatment effect: F(1,8)=7.16, p=0.028; interaction effect: F(12,96)=3.13, p=0.0009. **K:** Total average intersections for microglia. n=4-6 animals/group. Statistics: Unpaired t-test. t=2.68, p=0.028. **L:** Representative microglial images. Scale bar indicates 10 μm. For all statistics, *p<0.05, **p<0.01, ***p<0.001, ****p<0.0001. Error bars represent SEM.

We next investigated the extent of glial activation in our PTOA model. As expected from our clustering analysis in Figure 5, we observed an upregulation of GFAP, indicating the presence of astrogliosis (Figure 6G,I). While we did not observe an increase in total IBA1 intensity in the SDH (Figure 6H,I), Sholl analysis revealed a significant increase in the number of intersections in the PTOA dorsal horn, as well as an increase in the total number of branch intersections, suggestive of a phenotypic switch to a more ramified state with heightened activation (Figure 6J-L). These findings indicate convergent glial mechanisms for PTOA involving both astrocytes and microglia.

### 3.7 Inhibiting astrocytic gap junctions and microglia is analgesic in PTOA

Having confirmed that our PTOA model exhibited characteristics of astrogliosis and microgliosis in the SDH, we next sought to reverse these changes pharmacologically with the goal of inducing analgesia. As a proof of concept, we first tested a non-selective glial gap junction inhibitor, carbenoxolone (CBX), on our PAC pain model (Figure 7A-C).[19,60,80,93] Following an intrathecal injection of paclitaxel, we observed acute mechanical sensitization as early as thirty minutes (Figure 7A). Two hours after intrathecal injection of paclitaxel, an intrathecal injection of CBX was sufficient to transiently induce analgesia for at least three hours after intrathecal CBX injection (Figure 7B-C).

**Figure 7.**
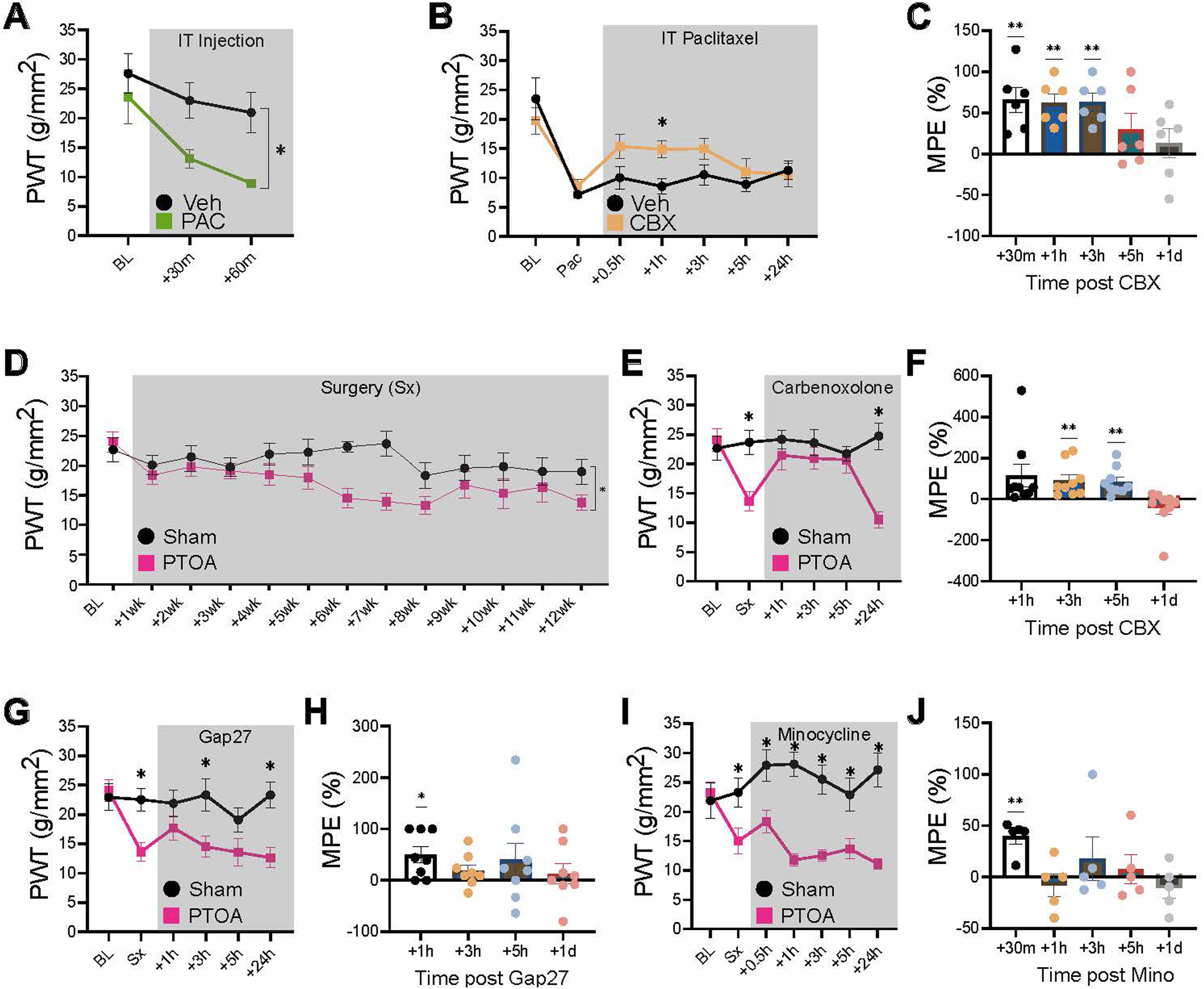
Inhibition of astrocytic gap junctions and microglia reverses post-traumatic osteoarthritis (PTOA) induced mechanical allodynia. **A** Time course of mechanical allodynia after an intrathecal injection of paclitaxel (50 nM, 4 μL). n=4 animals/group. Statistics: 2-way repeated measures ANOVA with Sidak’s post-hoc. Time effect: F(2,12)=9.62, p=0.0032; treatment effect: F(1,6)=7.078, p=0.038; interaction effect: F(2,12)=1.39, p=0.29. **B** Time course of analgesia following an intrathecal injection of carbenoxolone (5 μL, 50 nM) after intrathecal paclitaxel. n=4 animals/group. Statistics: Mixed-effects analysis. Time effect: F(2.986,29.36)=18.21, p<0.0001; treatment effect: F(1,10)=1.41, p=0.26; interaction effect: F(2.986,29.36)=2.94, p=0.049. **C** Analgesia is quantified by MPE. n=4 animals/group. Statistics: 1-sample t-test. **D** Time course of mechanical allodynia after Sham or PTOA surgery (Sx) over 12 weeks. n=12 animals/group. Statistics: 2-way repeated measures ANOVA with Sidak’s post-hoc. Time effect: F(12,143)=2.37, p=0.0082; treatment effect: F(1,143)=23.39, p<0.0001; interaction effect; F(12,143)=1.17, p=0.31. Nineteen weeks after surgery, an intrathecal injection of **E** carbenoxolone (5 μL, 50 nM) was injected. n=9-12 animals/group. Statistics: 2-way repeated measures ANOVA with Sidak’s post-hoc. Time effect: F(2.802,58.84)=4.79, p=0.0056; treatment effect: F(1,21)=6.32, p=0.020; interaction effect: F(2.802,58.84)=7.713, p=0.0003. **F** Analgesia is quantified by MPE. n=9-12 animals/group. Statistics: 1-sample t-test. **D** Following a week-long wash out period, the connexin mimetic peptide Gap27 (5 μL, 3 nmol) was intrathecally injected. n=8-12 animals/group. Statistics: 2-way repeated measures ANOVA with Sidak’s post-hoc. Time effect: F(5,100)=4.13, p=0.0019; treatment effect: F(1,20)=9.74, p=0.0054; interaction effect: F(5,100)=3.202, p=0.0101. **E** Analgesia is quantified by MPE. n=8-11 animals/group. Statistics: 1-sample t-test. **F** Following a week-long wash out period, the microglial inhibitor minocycline (5 μL, 100 μg) was intrathecally injected. n=5-9 animals/group. Statistics: 2-way repeated measures ANOVA with Sidak’s post-hoc. Time effect: F(6,78)=1.367, p=0.24; treatment effect: F(1,13)=77.26, p<0.0001; interaction effect: F(6,78)=3.487, p=0.0042. **G** Analgesia is quantified by MPE. n=5-9 animals/group. Statistics: 1-sample t-test. For all statistics, *p<0.05, **p<0.01, ***p<0.001, ****p<0.0001. Error bars represent SEM.

With evidence that suppressing astrocytic gap junctions during astrogliosis can be analgesic, we next asked if this approach could be used in PTOA to have a similar analgesic effect (Figure 7D-F). We observed a similar analgesic effect lasting for at least 3 hours after intrathecal injection of CBX in PTOA mice, compared to vehicle controls (Figure 7E-F). To isolate the effect of astrocytic gap junctions, we used a selective inhibitor, Gap27,[20,70,87,88] which demonstrated a significant, but more transient analgesic effect (Figure 7G-H). Since our LDA clustering analysis and Sholl analysis suggested that the PTOA model exhibited microglial activation, we next sought to inhibit microglial activation. Using minocycline, a microglial inhibitor,[29,42,96] we observed a transient reversal of mechanical allodynia at the thirty minute timepoint (Figure 7I-J). These pharmacological and behavioural results indicate concurrent contributions of both astrocytes and microglia to mechanical allodynia in PTOA.

## Discussion

A major challenge persisting in pain research is determining which chronic pain conditions involve central sensitization. Here, we developed a novel Ca^2+^ imaging and analysis pipeline to investigate central changes in rodent models of pathological pain. Using a supervised clustering approach, we discovered that PTOA exhibits an activity signature that overlaps with SNI and paclitaxel, both of which have well-documented central sensitization driven by glial activity. By targeting glial activity pharmacologically, we were able to observe analgesic effects.

Ca^2+^ imaging analysis pipelines have increased in functionality and accuracy alongside imaging advancements.[6,32,89] Here, our pipeline addresses holes specific to epifluorescent imaging, significant tissue drift and poor imaging quality, by adapting from two previous pipelines, Minian and MIN1PIPE.[17,59] One of the most used pipelines for Ca^2+^ imaging analysis, CaImAn, provides automated detection but requires expertise for customization.[27] Minian and MIN1PIPE offer automation options but have limited modularity.[17,59] CuMIN prioritizes modularity and real-time feedback over full automation, which we found more appropriate for videos with variable image quality, requiring manual ROI selection. Critically, our use of chemical Ca^2+^ indicators rather than genetically encoded sensors eliminates the need for transgenic animals, enabling application across species and potentially to human spinal cord tissue for translational studies. Future iterations could integrate computer vision -aided machine learning to automate analysis in challenging imaging conditions.[3,35]

During our experiments, we observed that some cells exhibited spontaneous events in the absence of stimulation, which we termed spontaneously active (SA), while another population was not spontaneously active but responded to glutamate, termed non-spontaneously active (NSA).

Spontaneous activity in SDH neurons has been documented both in *ex-vivo* slice preparations at over 60% and *in-vivo* approximately 5%.[18,79] We observed that in slices from SNI mice, activity became biased towards the SA population, suggesting enhanced cellular activity, recruitment to this population, or a combination. Spontaneous activity may reflect intrinsic homeostatic properties of certain neuronal populations, ongoing network activity, or both, and increased spontaneous activity has been associated with pathological pain states including central and peripheral neuropathic pain.[7,30,36,46]

The increased recruitment of NSA cells in response to glutamate following pre-treatment with bicuculline and strychnine is consistent with disinhibition of excitatory SDH neurons, as previous studies have established that chloride dysregulation preferentially unmasks excitatory neuron activity. [49,83] Under normal conditions, tonic inhibition generally suppresses neuronal activity, and when this inhibition is compromised, neurons that would normally remain silent become responsive to excitatory input.[15,39,83,99] Interestingly, we observed a similar phenotype for our CFA model as well, suggesting that disinhibition might also be present. This mechanism has been hinted at previously, where a peripheral CFA injection was observed to induce epigenetic downregulation of spinal KCC2 expression.[55,99] Our finding that spontaneous network activity (frequency and amplitude) was generally unchanged, while NSA recruitment increased, suggests that central sensitization primarily affects evoked rather than spontaneous activity patterns, at least at the timepoints examined.

There has been debate whether PTOA could be considered a model of nociceptive, neuropathic, or nociplastic pain.[12,81] Here, we leveraged supervised machine learning to answer this question, training an LDA on the rich quantitative output from Ca^2+^ imaging from four well-characterized models: CFA and capsaicin for nociceptive pain, SNI for lesion-induced neuropathic pain, and paclitaxel for chemotherapy-induced neuropathy. We then used this classifier to phenotype PTOA. As expected, capsaicin and CFA clustered together, with distinct clustering of SNI. Paclitaxel’s intermediate position with partial overlap with CFA was consistent with documented astrocytic activation.[11,25,38,57,98,100] Notably, PTOA showed a scattered distribution with significant paclitaxel and SNI overlap, suggesting that the underlying mechanisms of PTOA might have shared characteristics of both. Molecular validation confirmed that PTOA samples exhibited robust astrogliosis, matching our paclitaxel samples, as well as microgliosis, quantified with Sholl analysis. Pharmacological targeting of astrocytic gap junctions using carbenoxolone and Gap27 and microglial activity using minocycline produced analgesia, confirming that microgliosis and astrogliosis indeed are contributing to mechanical allodynia. This approach transforms Ca^2+^ imaging from a descriptive tool into a platform for mechanism discovery and drug screening, where compounds can be tested for their ability to normalize pain-associated signatures toward naive patterns.

Our findings are consistent with an emerging body of evidence that osteoarthritis involves central sensitization mediated by spinal glia. Previous studies using MIA-induced osteoarthritis models have demonstrated increased microglial reactivity coincident with mechanical hyperalgesia, which could be attenuated by intrathecal minocycline or fluorocitrate.[1,64] In MIA models, microglial P2X7 receptor signaling and pannexin-1 channels have been implicated in mechanical hypersensitivity, and ablating microglia using Mac1-saporin attenuates hyperalgesia.[53,64] Recent studies using more traumatic models of OA, such as a partial medial meniscectomy, have characterized evidence of microgliosis and astrogliosis.[28,61,62,74] The mixed phenotype we observed in PTOA, sharing features with both paclitaxel and SNI, aligns with clinical observations that osteoarthritis pain often persists despite minimal joint inflammation, suggesting significant central contributions.[4,58]

The analgesic efficacy of astrocytic gap junction inhibitors in PTOA is noteworthy, as connexin - 43-mediated signaling has been implicated in the maintenance of late-phase neuropathic pain through astrocytic chemokine release.[11,25,100] Intrathecal injections of carbenoxolone and selective connexin-43 blockers (Gap26 and Gap27) reduced mechanical allodynia when administered weeks after nerve injury, suggesting these interventions may be effective even in established pain states.[11,94]

Importantly, carbenoxolone and Gap27 have shown efficacy in multiple pain models, including sciatic nerve injury,[77] spinal cord injury,[71] and cancer-induced bone pain,[21] suggesting that astrocytic gap junction signaling may represent a conserved mechanism across diverse pain conditions and a promising target for PTOA pain management.

A consideration when interpreting our data is the cellular source of Ca^2+^ signals. As our imaging approach did not filter for cell type, the Ca^2+^ signaling signatures we observed likely reflects changes in heterogeneous populations, including non-neuronal cells such as astrocytes.[38,76,95] Astrocytic Ca^2+^ responses can arise from nerve injury, and directly activating spinal astrocytes can induce mechanical and thermal hyperalgesia in uninjured animals.[37,40] Cell type-specific genetically encoded Ca^2+^ indicators could resolve this ambiguity,[2,34] but restricting detection to single cell types may undermine CuMIN’s ability to identify broad changes in network activity, as the interaction among neurons, astrocytes, and microglia appears central to pain pathophysiology. [38] An alternative approach is to wash on glial specific fluorescent markers, such as SR101 for astrocytes or CDr20 for microglia.[50,66]

Ca^2+^ signaling in SDH neurons can arise from extracellular and intracellular sources, including voltage-gated Ca^2+^ channel activation during action potential firing, Ca^2+^-permeable AMPA receptor activation, NMDA receptor-dependent Ca^2+^ influx, metabotropic glutamate receptor-triggered release from intracellular stores, and Ca^2+^-induced Ca^2+^ release from the endoplasmic reticulum.[8,13,32,33,44,48,52,90] In neuropathic pain conditions, increased activation of T-type Ca^2+^ channels has been observed to drive nociceptor hyperexcitability.[33,48] Similarly, CP-AMPARs are upregulated in superficial SDH neurons following nerve injury and inflammatory pain and contribute to enhanced Ca^2+^ influx.[22,43,44] Doolen et al. (2012) demonstrated that glutamate-evoked Ca^2+^ events were larger on the side ipsilateral to nerve injury, indicative of increased glutamate receptor density or conductance.[18] NMDA receptor phosphorylation is also enhanced in pain states and contributes to central sensitization.[10,52] The precise contributions of each mechanism to the signatures we observed remains unknown, but the association between Ca^2+^ signature and analgesic pharmacology suggests the observed Ca^2+^ signatures capture conserved and functionally relevant network changes.

Our approach has several limitations that should be acknowledged. First, all experiments were performed with male mice. It is therefore of particular importance to confirm in future experiments that microgliosis and astrogliosis occur in female post-traumatic osteoarthritis mice as well. With respect to mechanistic limitations, the *ex-vivo* preparation lacks descending modulatory input from brainstem nuclei, which exert top-down control over spinal nociceptive processing, as well as the peripheral input from primary afferents and sensory neurons. The transverse slice preparation, while enabling optical access across local ascending and descending networks, disrupts lateral connectivity. Tissue preparation effects may alter baseline network activity compared to *in-vivo* conditions, where spontaneous Ca^2+^ events occur at higher frequencies.[75] Additionally, while bulk-loaded Ca^2+^ indicators provide excellent signal-to-noise ratios, they do not permit genetic targeting of specific cellular populations. Of note, the timepoints chosen are a snapshot representative of the behavioural readout and are typical for mechanistic studies involving these pain models. Whether the changes we observe for SNI and PTOA were present at earlier timepoints or persist weeks after is uncertain. Nonetheless, our approach can be used to study long-term central changes in these pain models, particularly the transition from acute to chronic pain.

Overall, this study establishes Ca^2+^ imaging as a viable approach for phenotyping central sensitization across pain conditions and challenges our current understanding of PTOA as simply a model of peripheral inflammation. Using CuMIN, we demonstrate that PTOA, a clinically relevant model of osteoarthritis, exhibits a mixed central sensitization phenotype with features of both chemotherapy-induced neuropathy and nerve injury. This finding has immediate therapeutic implications that support targeting spinal glia rather than treating osteoarthritis solely as a peripheral inflammatory condition. The analgesic efficacy of carbenoxolone, Gap27, and minocycline in PTOA validates this mechanistic insight and identifies astrocytic gap junctions and microglial signaling as druggable targets for osteoarthritis pain. Collectively, our results demonstrate that CuMIN provides a framework for mechanism discovery and phenotype-guided drug development in chronic pain.

## Acknowledgements

We thank the members of the Bonin lab for their critical appraisal of this manuscript. Claude Opus 4.5 assisted in the development and troubleshooting of CuMIN.

## Author contributions

S.W.F., E.K.H., J.B., J.R.M, M.E.H., J.A.S., and R.P.B. contributed to the project design. S.W.F., E.K.H., H.Z., and J.A.S. conducted experiments. S.W.F., E.K.H., J.K.C., J.L.T.D., S.M.N, and J.A.S analyzed data. S.W.F. created CuMIN. S.W.F., J.K.C., and J.L.T.D performed technical validation. S.W.F., J.K.C., J.L.T.D., and J.A.S. performed and analyzed immunofluorescence data. S.W.F. performed behavioural testing. S.W.F., E.K.H., and R.P.B. wrote the manuscript, and all authors edited and approved the final manuscript.

## Data and materials availability

All data needed to evaluate the conclusions in the paper are present in the paper and/or the Supplementary Materials.

